# Gabija restricts phages that antagonize a conserved host DNA repair complex

**DOI:** 10.1101/2025.08.30.673261

**Authors:** Alex Hong, Miaoxi Liu, Alexis Truta, Alexander Talaie, Gerald R. Smith, Joseph Bondy-Denomy

## Abstract

Anti-bacteriophage systems like restriction-modification and CRISPR-Cas have DNA substrate specificity mechanisms that enable identification of invaders. How Gabija, a highly prevalent nuclease-helicase anti-phage system, executes self-vs. non-self-discrimination remains unknown. Here, we propose that phage-encoded DNA end-binding proteins that antagonize host RecBCD sensitize phages to Gabija. When targeting a temperate Lambda-like phage in *Pseudomonas aeruginosa*, Gabija protects the cell by preventing phage genome circularization and subsequent replication. Phage and plasmid sensitivity to Gabija is licensed by DNA end-binding complexes such as a phage exonuclease together with a ssDNA-annealing protein or Gam_Mu_ dimers, which prevent loading of host repair complex RecBCD. Escape phages lacking these end-binding proteins become protected from Gabija by RecBCD translocation activities. RecBCD activity on the bacterial genome also prevents Gabija from targeting self-DNA. Therefore, we propose that Gabija antagonizes circularization of linear DNA devoid of RecBCD as a mechanism to identify foreign invaders.

## Introduction

The bacteria-phage arms race has driven the generation of an incredible range of bacterial antiphage immune systems that identify foreign invasion and prevent phage replication. To date, the prokaryotic anti-phage defense repertoire is predominated by DNA-targeting systems with a large portion of this arsenal populated by CRISPR-Cas and restriction modification (RM) systems^1,2^. However, the recent flourish in discovery of new bacterial immune systems has revealed many unique DNA-targeting systems.

Of these systems, Gabija is one of the most common, identified in 13.13% of sequenced prokaryotic genomes, and encoded in host genomes, mobile islands, and plasmids^1,3–6^. Gabija is one of the 6 most common systems across prokaryotes and within strains of the opportunistic generalist human pathogen, *Pseudomonas aeruginosa* (Pa)^1,2^. Gabija is a two gene nuclease-helicase system first identified by Doron, Melamed et al^3^. Both of Gabija’s genes - GajA, an OLD-like (overcoming lysogenization defect) nuclease, and GajB, a UvrD-like helicase - are required for phage restriction (Figure 1A)^7–10^. GajA contains an ATPase-like domain and a topoisomerase-primase (TOPRIM) domain, the latter of which is believed to act as an endonuclease or exonuclease depending on its substrate^8^. The OLD nuclease was first defined as a single-gene P2 prophage-encoded system that prevented λ phage superinfection. It was later discovered that λ phages escape P2 OLD upon deleting their red recombination system (*exo*, *beta*, and *gam_λ_*)^7,11,12^. The TOPRIM domain characteristic of OLD enzymes has recently also been described in Retrons and the Ocr-triggered defense system, PARIS^13,14^.

**Figure 1.**
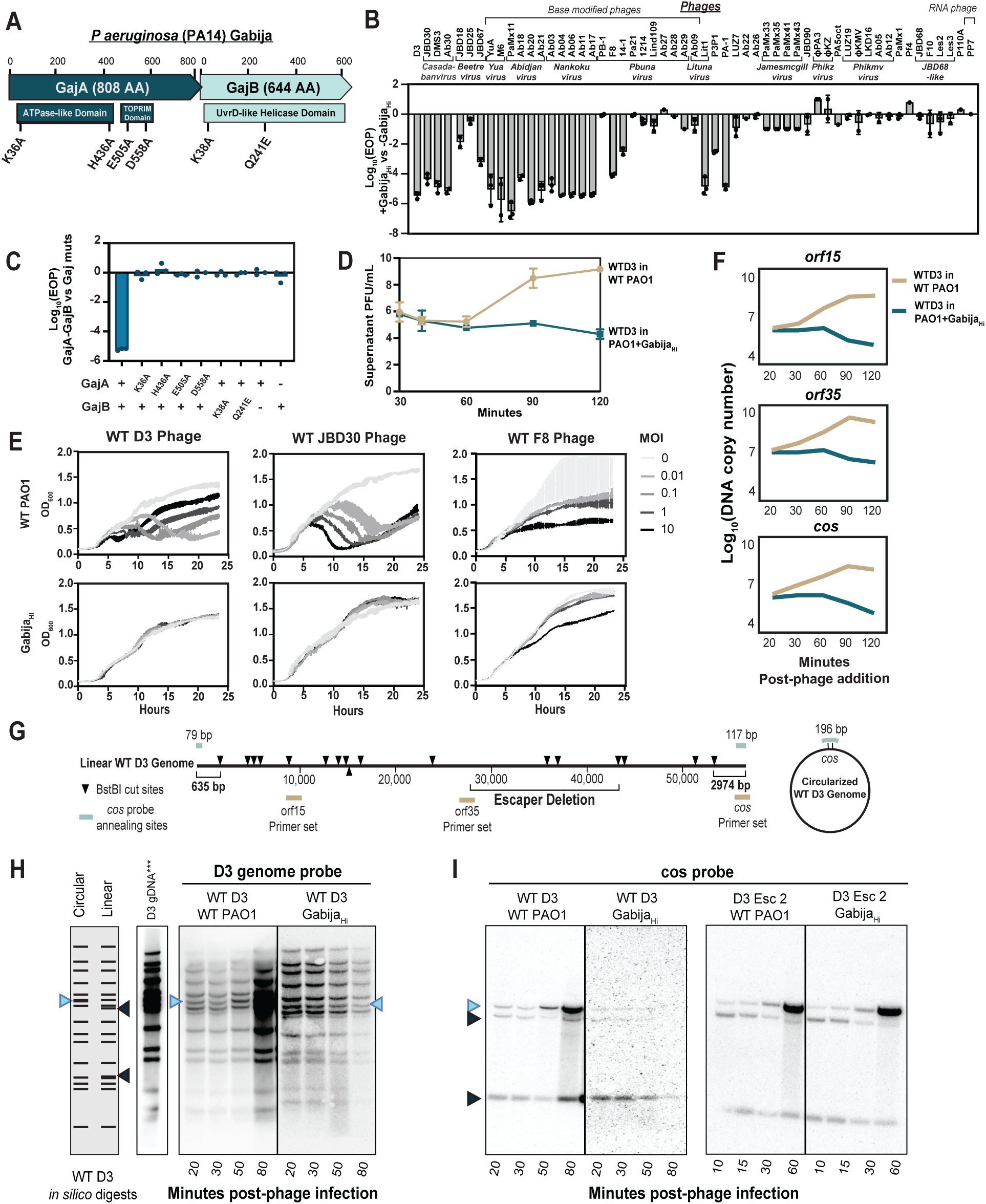
Gabija_PA14_ is a generalist non-abortive DNA-targeting system that blocks circularization of the D3 phage genome A. The PA14 Gabija operon B. Efficiency of plating (EOP) was quantified by the amount of plaque-forming units PFUs/mL on PAO1 + Gabija_Hi_ divided by PFU/mL on WT PAO1 (n = 3). Data are mean ± SD. Phages are grouped by genera in brackets. C. EOP of D3 phage on strains with GajA-GajB point mutations or GajA/GajB only. D. Extracellular D3 titer during a single step infection of PAO1 +/- Gabija. D3 phage was allowed to adsorb with Pa strains at room temp or on ice for 20 minutes prior to wash steps. Supernatant samples were then collected (30 min = post-wash) and PFU was calculated by spot titration on naive PAO1 lawns (n=3). Data are mean ± SD. E. Growth curves (OD_600_ measurements) of phages D3, JBD30, and F8 without (WT PAO1) or with Gabija_Hi_ at multiplicity of infections (MOIs) of 0.01, 0.1, 1 and 10 over 24 hours (n=3). Data are mean ± SD. F. Relative copy number of D3 phage genomic loci over time. G. A D3 phage genomic representation of the qPCR and Southern blotting analyses. qPCR probed regions are marked with yellow bars. BstbI cuts sites (black arrows) and the expected annealing sites of the *cos* probe (blue bars) are marked according to Southern blot analysis (Figure 1H and 1I). H. Southern blot analysis to query the integrity of the D3 phage genome over time with and without active Gabija. D3 genomic DNA was detected using P^32^ dCTP labelled probes generated either by random hexamer extension on D3 gDNA or *cos*-specific primed extension (n=3). An *in silico* digest of either circularized or *cos*-end linearized D3 genomic DNA is represented on the left. WT D3 genomic DNA was used as a control. I. Southern blot of the same membranes from 1H stripped and reprobed with a cos-specific probe (n=3). Black arrows indicate the two linearized ends of the phage genome. Blue arrows indicate the two linear ends coming together to form the larger band.

DNA-targeting systems often possess substrate specificity requirements that allow them to distinguish between self- and non-self-DNA. For example, innate immune systems like restriction modification (RM) systems detect the presence or absence of DNA modifications while adaptive CRISPR-Cas systems utilize sequence-specific guides to target defined nucleotide sequences. Ideally, these parameters ensure that the host DNA is not damaged following activation of these systems. These specificity requirements established RM and CRISPR-Cas as revolutionary biological research and genetic engineering tools. As DNA-targeting systems, the Gabija family likely also has substrate specificity requirements that ensure protection from self-targeting; however, this has not been investigated.

Previous work on Gabija has explored systems from *B. cereus* strain VD045 (Gabija_VD045_) and *E. coli* strain ECOR94^6,15–18^. *In vitro* cleavage assays suggested that Gabija_VD045_ is inhibited by high concentrations of NTPs and dNTPs, leading Cheng et al^17^ to propose that Gabija becomes an activated promiscuous nuclease after sensing nucleotide pool depletion during phage replication and transcription. Here, we characterize Gabija’s anti-phage mechanisms within *P. aeruginosa*. We show that *P. aeruginosa* strain PA14’s GajA-GajB (Gabija_PA14_) system is non-abortive and prevents phage replication initiation by targeting phage genome ends and preventing phage genome circularization. Therefore, Gabija is active prior to the initiation of any replication or transcription event. Phages can harbor one or more proteins that license Gabija targeting, each with unique mechanisms that hinge on the status of the host’s DNA repair/recombination system, RecBCD. We show that prevention of RecBCD loading on phage genomes licenses Gabija activity. Moreover, Gabija resistant phages, escape phages, and the host can become Gabija sensitive upon genetic ablation of RecBCD. This suggests that Gabija uses DNA that is devoid of RecBCD activity as its determinant for specificity to antagonize the circularization of linear DNA. In other words, RecBCD protects linear DNA from Gabija. Together, we demonstrate that Gabija_PA14_ prevents end joining of linear DNA that is unbound by RecBCD *in vivo*, and that phages often prevent RecBCD loading, which sensitizes them to Gabija.

## Results

### Gabija_PA14_ is a non-abortive generalist antiphage system

To establish a model system for Gabija, we identified an intact *gajA-gajB* operon in model strain PA14 within the PAPI-1 pathogenicity island, a 115-gene conjugative element key for many virulence phenotypes^19^. GajA, GajB, or the full GajA-GajB system were expressed in the phage-sensitive strain, PAO1, which lacks a functional Gabija system and has high degree of genetic conservation with PA14, barring PA14’s two pathogenicity islands^19,20^. PA14 GajA and GajB were both required for broad antiphage defense and are, together, most optimal at 30 °C (Figure 1B and S1). Point mutations in GajA’s ATPase and topoisomerase-primase (TOPRIM) domains and GajB’s ATPase and DExx (DESQ) motif active sites confirm that conserved residues are required for antiphage defense (Figure 1C). These characteristics are consistent with previous findings using *Bacillus cereus* Gabija^15–18^.

We then established two PAO1 chromosomally inserted Gabija systems that will be used for the rest of this study. These either included the 250 bp native upstream sequence found in the PA14 genome (Gabija_Lo_) or excluded the native upstream sequence (Gabija_Hi_). Interestingly, each of these strains has different levels of anti-phage restriction across our PAO1 phage panel. Gabija_Hi_ had the broadest and strongest anti-phage activity and was constitutively active as it did not require, nor did it improve upon, induction via the co-inserted *lac* promoter. Gabija_Lo_ only restricted phage upon IPTG induction and had reduced phage restriction across fewer phages, suggesting that the native upstream sequence is repressive under these growth conditions. Both Gabija_Hi_ and Gabija_Lo_ target an extensive panel of phages, including unrelated phage families, suggesting that Gabija has a broad ability to antagonize phage.

To determine if Gabija_PA14_ is abortive, we measured OD_600_ of PAO1 cultures with or without Gabija_Hi_ infected by either temperate phages D3 (λ-like, *Detrevirus*) or JBD30 (Mu-like, *Casadabanvirus*), or lytic myophage F8 (*Pbunavirus*) with multiplicities of infection (MOIs) from 0.01 to 10 (Figure 1E). Gabija_Hi_ was observed to robustly prevent lysis of PAO1 by all three phages in almost all MOI conditions, with the exception of F8 phage at an MOI of 10. Interestingly, we did not find any population collapse or dormancy in any of the above conditions, suggesting that Gabija_PA14_ is not only non-abortive, but that it can likely directly antagonize unrelated phages. These results contradict the previously presented data that Gabija is an abortive infection system^17^.

### Gabija blocks circularization of D3 phage genome

To determine when during the phage infection cycle Gabija induces phage failure, we executed a single step infection with phage D3. During D3 infection of PAO1 at an MOI of 1 at 30 °C, phage-induced bacterial lysis occurs between 60-90 minutes in the absence of Gabija (Figure 1D). In the presence of Gabija, no phage production is detected. Phage DNA replication in the Gabija_Hi_ strain was measured by qPCR of three loci (across cohesive *cos* ends or within *orf15* or *orf35)* at 20, 30, 60, 90, and 120 minutes post-phage introduction. For all three loci, copy numbers increase by 2-3 orders of magnitude by 60 minutes in the absence of Gabija_Hi_. Genome replication is not observed in the presence of Gabija_Hi_, where DNA copy number decreases by 2 orders of magnitude 60 minutes post-adsorption (Figure 1F and 1G). This suggests that Gabija is antagonizing DNA replication or a prior step. Note that the decrease in phage gene copy number at later time-points is relative to host DNA which is likely increasing during this time frame. Therefore, whether Gabija is actively degrading phage DNA is not yet clear.

To assay both the timing of Gabija’s DNA-targeting activity and the effects of its activity on the phage gDNA at a higher resolution, Southern blotting was performed on gDNA extracted from liquid infections of D3 phage (Figure 1G and 1H). The D3 genome was visualized by digesting extracted total lysate gDNA with the BstBI restriction enzyme, then probed with D3 gDNA generated P^32^-labeled probes. In WT PAO1, the entire WT D3 genome gains signal intensity between 50-80 minutes - suggesting that this is when DNA replication occurs - then further increases in intensity at 80+ minutes. At the 50-minute time point, the band that indicates the cohesive (*cos*) ends coming together - creating a circular genome - becomes more intense relative to the other bands of the genome at the same time point. Alternatively, in the presence of Gabija_Hi_, the entire WT D3 phage genome decreases in concentration over time but with no clear cleavage event or specific band disappearing relative to others. Interestingly, we noticed that the band corresponding to circularized genomes never appears in the presence of Gabija, and the relative intensity of all bands stay the same at each time point, suggesting that circularization of the phage genome never occurs. To confirm this observation, we stripped the membranes and re-hybridized with new probes specific to the D3 *cos* ends (Figure 1I). Indeed, without Gabija present, we see circularization of phage gDNA (an increase in intensity of the top circularized band accompanied by a decrease in intensity of the two lower bands that indicate linearized genome ends) accompanied by replication and subsequent *cos* cleavage during packaging. But in the presence of Gabija, the circularized upper band does not grow in intensity, suggesting that the phage genomes do not circularize and, subsequently, do not replicate. Together, these data demonstrate early phage failure and suggest that Gabija has a tropism for interfering with genomic ends.

### Phage restriction by Gabija is licensed by a putative D3 recombination system

To understand the phage factors that license Gabija targeting, we isolated spontaneous D3 phage escape mutants in the presence of Gabija, with an average escape plaque frequency of 3.9 x 10^-5^. Of 43 independently isolated escape phages, we identified 8 unique escape genotypes with deletions ranging from 6.7-16.9 kb, all centered around a minimal deletion of genes *orf52*-*orf66* (Figure 2A). All D3 escape phages replicate just as well in WT PAO1 as in PAO1 + Gabija_Hi_ demonstrating complete Gabija escape. Remarkably, these deletions are well-tolerated and do not obviously hinder maximum phage replicative titer, except in the presence of a CRISPR-Cas system (see below). D3 escape phage 4 (Esc 4) has the smallest deletion while Esc 2 had the largest deletion that spanned *orf35* – its integrase – to *orf67*. The loss of *orf35* converted escaper 2 to a clear-plaquing virulent mutant unable to form lysogens. When probed via the Southern blotting experiment presented above, D3 Escape mutant 2 replicated efficiently and circularized rapidly, even in the presence of Gabija (Figure 1I). Interestingly, D3 Esc 2 appears to circularize at an earlier time point compared to WT D3 when assessed with a *cos*-specific Southern probe.

**Figure 2.**
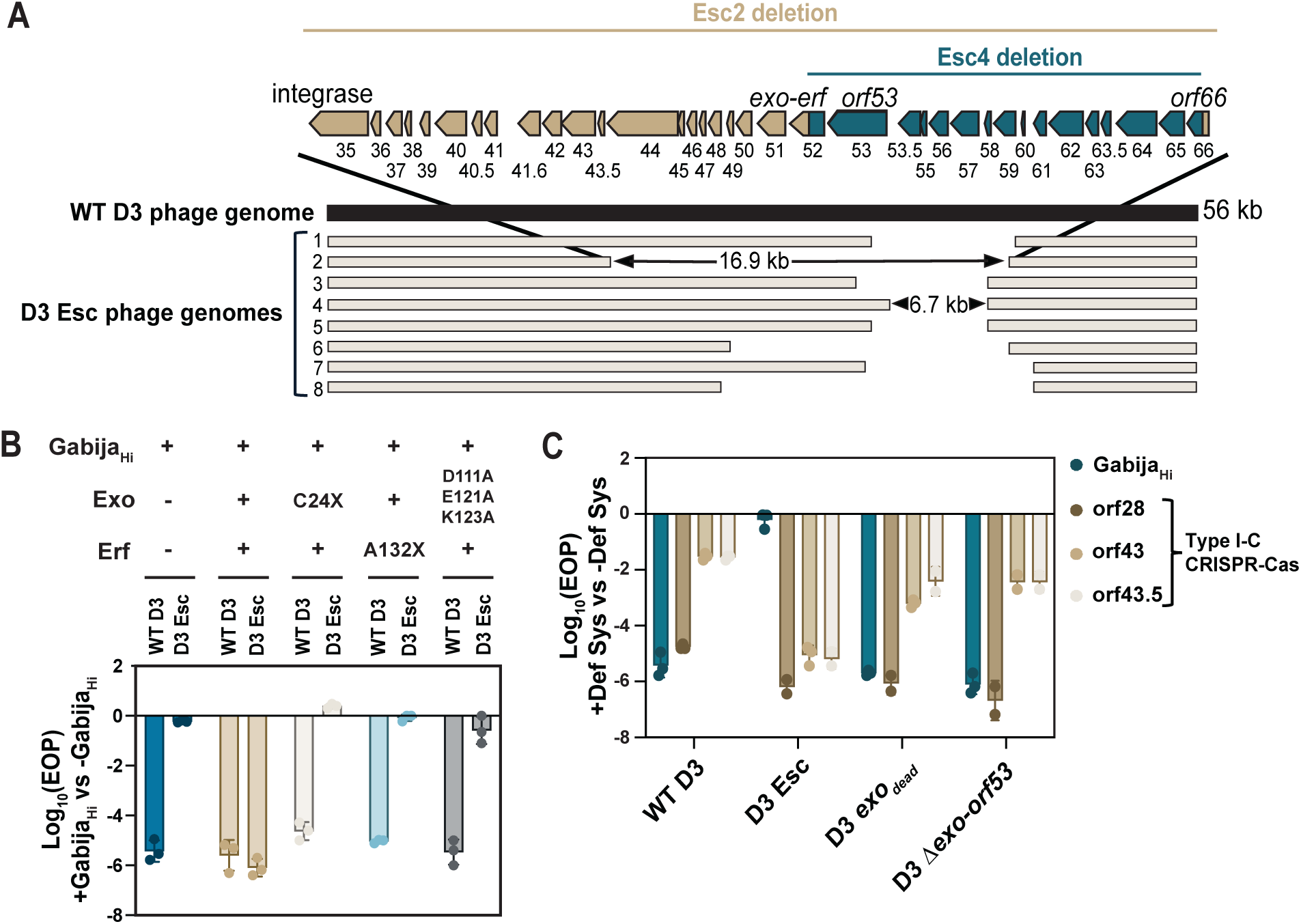
D3 phage deletes the Exo-Erf system to escape Gabija targeting A. Schematic of WT D3 (dark navy) and D3 escape phage (light tan) genomes. The largest Esc 2 (tan) and smallest Esc 4 (blue) deleted genes are represented above. Table S1 has records of putative protein functions based on homology searches. B. Efficiency of plaquing of WT D3 and D3 Esc phages with overexpression (OE) of Exo-Erf, truncated Exo or truncated Erf, or catalytically dead triple mutant Exo (n=3). EOPs were calculated by dividing PFU/mL on Gabija + OE of the complemented genes divided by PFU/mL on PAO1 with OE of the respective genes. C. Efficiency of plaquing of D3 variants in the presence of either Gabija or Type I-C CRISPR-Cas targeting one of three D3 genes.

We hypothesized that Gabija either targets DNA sequence(s) within the deleted region or is “activated” by one or more of the genes deleted in the escaper phages. We first queried Esc 4 as it has the smallest deletion. We engineered recombinant D3 Esc 4 phages by reinserting 1100 bp sections of the 6.7 kb deleted region back into Esc 4’s deletion junction to determine if any sequences restored Gabija targeting (Figure S2). In parallel, we expressed each of the 16 genes from Esc 4’s deleted region in Gabija_Hi_ to determine if any restore targeting of D3 escape phages *in trans*. Expression of gp52 *in trans* restored targeting of D3 Esc 4, as did restoration of this region in the genome (Figure S3). Interestingly, gp52 expression did not restore targeting of any other D3 escape phages, suggesting that an additional missing gene from these phages is required to sensitize phages to Gabija. Expression of gp51, the protein encoded by the neighboring gene, together with gp52 *in trans* led to targeting of all escaper phages (Figure 2B). Homology searches revealed that gp51 is a homolog of the exonuclease (Exo) from λ phage’s Red recombination system while gp52 is a single strand annealing protein (SSAP) of the essential recombination function (Erf) family, first discovered in phage P2^21–26^. Together, these form a putative recombination system analogous to the λ Red recombination system. Early stop codons in Exo or Erf, or a triple point mutation in the catalytic domain of Exo, eliminated complementation of escape phage targeting, demonstrating that Exo and Erf function together to sensitize D3 phage to Gabija.

### D3 phage trades-off Gabija evasion with CRISPR-Cas sensitivity

Repair systems like the Red recombination system help evade Type I and Type II CRISPR-Cas targeting^27–29^. We therefore asked if Gabija escape, achieved through losing a putative recombination system and flanking genes, led to a trade-off with increased sensitivity to a *P. aeruginosa* CRISPR-Cas system. Type I-C CRISPR-Cas targeting of essential gene *orf28* led to restriction of plaquing by 5-6 orders of magnitude across all WT and escape D3 phages (Figure 2C). However, crRNA sequences against non-essential regions (*orf43* and *orf43.5*) targeted D3 Gabija escapers 1-4 orders of magnitude more efficiently than WT D3 phage. A catalytically dead double point mutation in *exo* (exo_dead_) makes D3 phage 1-2 orders of magnitude more CRISPR-Cas sensitive than wild-type D3, depending on the crRNA used. This not only suggests that *exo-erf* is a functional recombination/repair system, but additionally demonstrates that other D3 genes missing in the escape deletion also promote CRISPR evasion or DNA repair. Therefore, D3 phage is sensitized to Gabija by a DNA recombination/repair locus that antagonizes CRISPR targeting. This generates a mechanistic opposition between two common DNA-targeting immune systems, forcing a trade-off.

### Gabija specifically favors injected DNA ends

Upon adsorbing to their hosts, dsDNA phages inject linearized genomes, which circularize before proceeding to either lytic growth or lysogeny. As Southern blotting revealed Gabija’s antagonism of genomic end circularization, we next tested whether these linear ends, together with Exo-Erf, are necessary for Gabija sensitivity. Given that a prophage will excise directly into its circular form and directly enter the lytic cycle (Figure 3A), we hypothesized that this would effectively bypass Gabija, despite undergoing identical DNA replication and packaging (two other moments where linear phage DNA can be exposed). D3 lysogens were isolated in cells lacking Gabija and then transformed with a plasmid-expressed Gabija (Gabija_30T_) or empty vector. For negative targeting controls, we also isolated D3 escaper phage lysogens. We first confirmed that Gabija_30T_ in D3 lysogens is still functional as unrelated phages JBD30 and obligately lytic YuA were still well-restricted by Gabija (Fig 3B). We then asked if Gabija targets the prophage or inducing prophage. Gabija_30T_ targeting of a sequence in the D3 prophage would result in DNA degradation of the host genome and, consequently, host death. However, D3 lysogens both without or with induced Gabija_30T_ do not exhibit any growth defect when measuring OD_600_ over time, with similar growth trends as WT PAO1 (Figure 3C). This demonstrates that a simple sequence motif is unlikely to be sufficient to induce targeting, the way a CRISPR-Cas system would work, for example.

**Figure 3.**
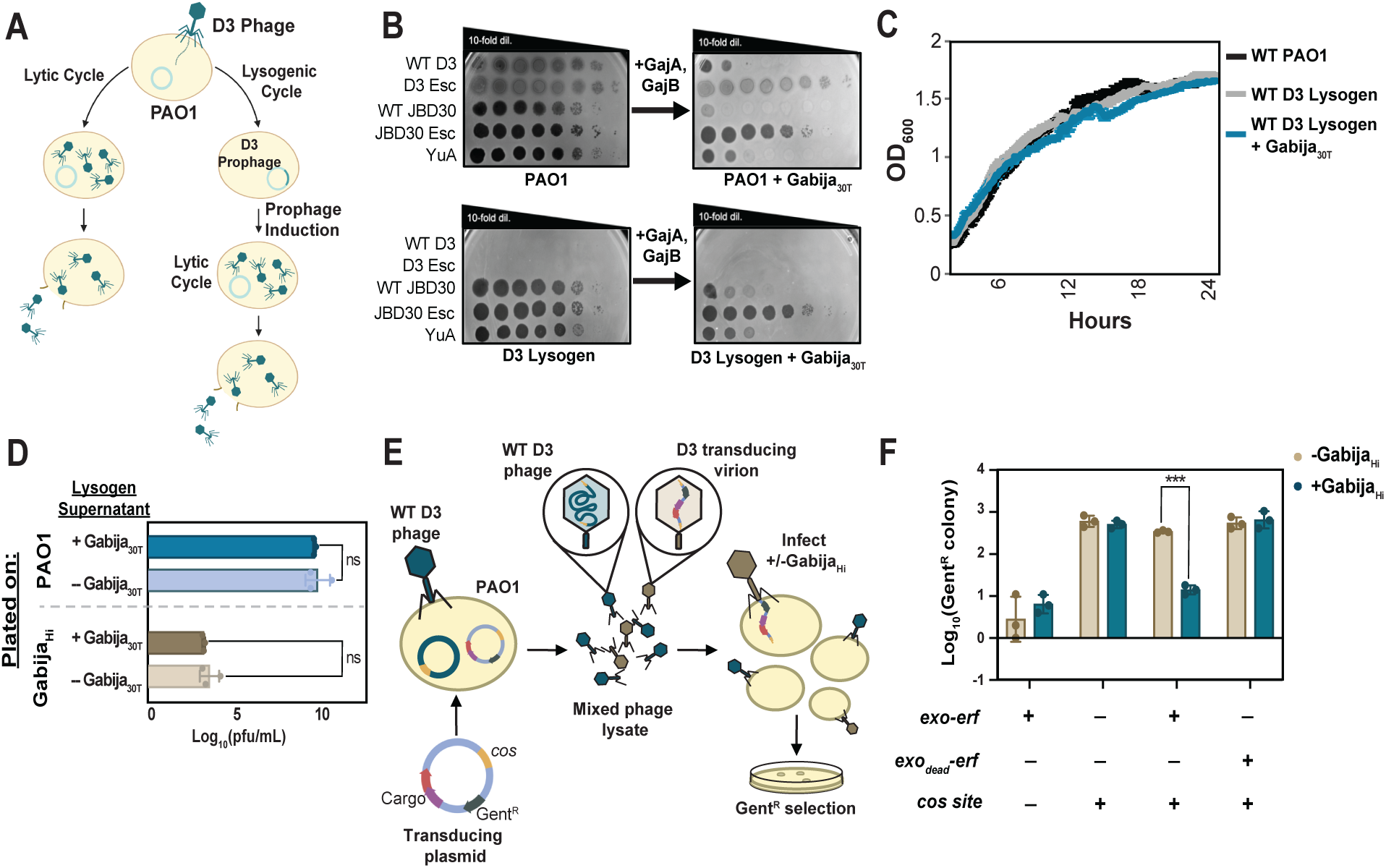
Gabija targets exposed ends of dsDNA A. The temperate phage life cycle. B. Plaque assays on WT PAO1, PAO1 + Gabija_30T_, WT D3 lysogen, and WT D3 lysogen + Gabija_30T_. C. Growth curves (OD_600_ measurements) of PAO1 cells with or without D3 prophage and Gabija_30T_ (n=3). Data are mean ± SD. D. Log-scale titer (PFU/mL) of phage from the supernatant of induced lysogens with or without active Gabija. These were titered on either a naive WT PAO1 lawn (blue bars) or a naive Gabija_Hi_ lawn (brown bars) (n=3). Data are mean ± SD. E. A schematic of the D3 transduction assay. Plasmids harboring the D3 *cos* sequence and gentamicin resistance +/- phage genes were transformed into PAO1. D3 lysates grown from these *cos*-plasmid containing cells would have a mixed population of WT D3 (blue) or transducing virions (brown). Successful transduction of the *cos* plasmid was calculated by colony count on gentamicin selection plates. F. Transduction efficiency (log-scale of Gent^R^ colony counts) with (blue) or without (tan) Gabija (n=3). Data are mean ± SD.

To test whether DNA replication or packaging are directly antagonized by Gabija, we next assessed the fate of the inducing prophage by measuring the phage titer in the supernatant. D3 prophages spontaneously induce, so the supernatant of overnight lysogen cultures +/- Gabija were collected, then titrated on naive WT PAO1. The titer of induced prophage did not change when Gabija was present (Figure 3D), indicating that Gabija activity does not limit phage success during prophage excision, phage DNA replication, or packaging. To check if these findings were misled by the presence of genetic mutants or epigenetic escaper phages that form prior to or during prophage establishment, the same induced phages were titrated on a naive Gabija_30T_ strain (no lysogen). We found that the induced phages remain fully sensitive to Gabija upon initiating a new infection, suggesting that these phages are genetically unchanged and that any epigenetic modifications acquired during lysogeny do not interfere with Gabija targeting.

Together, these results demonstrate that Gabija does not target D3 prophage nor a phage that is replicating in the lytic cycle that has derived from a prophage. This suggests that it is necessary for Gabija to be active against a phage in its initial linear form prior to prophage establishment or the induction of replication. Since DNA replication, repair intermediates, and nucleotide depletion will all be similar whether the phage replicates from an injected form or an induced prophage, we can rule out those events as playing a key role in the Gabija mechanism. We therefore propose that the exposed ends of injecting phage genomes, along with the Exo-Erf end-modifying proteins present a favored substrate for Gabija.

We next test this model and directly query whether linear ends, along with Exo-Erf, are sufficient to license Gabija against a DNA substrate. We took advantage of D3’s ability to transduce plasmids containing its 9 bp *cos* (5’-gcgccccca-3’) sequence^30^. The *cos* sequence is cleaved by the phage terminase during packaging to form cohesive sticky ends, which dictates the ends of the linearized phage genome. A basic cloning vector with a Gent^R^ marker was modified to possess a *cos* site flanked by its putative terminase-binding sequences, and to express Exo-Erf from a pBAD promoter (Figure 3E). D3 phage was grown on PAO1 possessing these *cos* plasmids to generate transducing phage lysates where some virions are packaged with plasmids, likely as concatemers. The lysates were mixed with cells with or without Gabija and transduced Gent^R^ colonies were counted.

In the absence of a *cos* site on the plasmid, no transduction was detected (Figure 3F). *cos*-only plasmids were well transduced, but we saw no significant difference in colony counts between cells with or without Gabija, demonstrating that Gabija does not generically target injected phage/plasmid DNA ends. Gent^R^ counts significantly decreased only upon transduction of plasmids harboring both Exo-Erf and *cos* in a Gabija-dependent manner, demonstrating that *cos* and Exo-Erf are sufficient to sensitize a DNA substrate to Gabija. *cos* plasmids with a catalytically dead Exo and Erf are transduced at similar rates with and without Gabija, suggesting that Exo’s resection activity on the DNA, which likely creates a substrate for Erf binding, generates a Gabija substrate. Together, we show that it is necessary for Gabija to act early in D3 phage infection, that DNA ends and an end-functioning DNA-recombination system are sufficient to sensitize a DNA substrate, and that the action of Gabija prevents phage genome circularization (Figure 1I) and therefore, DNA replication.

### Gabija licensed by Mu Gam, a distinct end-binding protein family

Despite the clear demonstration of the importance of Exo-Erf in licensing targeting of phage D3, two results pertaining to D3 remained perplexing. First, the D3 escape phages all had deletions of >16 genes in addition to *exo-erf*, for reasons that were not clear. Second, we found that an engineered D3 *exo*_dead_ phage does not escape Gabija (Figure 4A). Therefore, Exo-Erf is sufficient to induce Gabija targeting of a phage or plasmid but in the WT phage background, there may be redundant factors that license Gabija sensitivity. To identify these genes, we attempted to complement the escape phenotype (i.e. reinstate Gabija targeting) with genes *orf53-66*. These experiments revealed that *orf53* is also independently a licensing factor of Gabija, but *orf53.5-65* are not (Fig S2). In addition, expression of *orf66*, which falls at the other border of the minimal escape phage deletion, was toxic to the host in a Gabija-dependent manner. That is, neither Gabija_Hi_ nor Gabija_Lo_ strains were viable upon *orf66* expression, but *orf66* was not toxic in a strain lacking Gabija. This suggested that *orf66* over-expression may spuriously induce Gabija activity towards self. Regardless, these observations reconcile the fact that all escape phages require, at minimum, the 6.7 kb deletion that removes all three D3 genes that can license Gabija targeting – *exo-erf* (*orf51-52*), *orf53*, and *orf66*.

**Figure 4.**
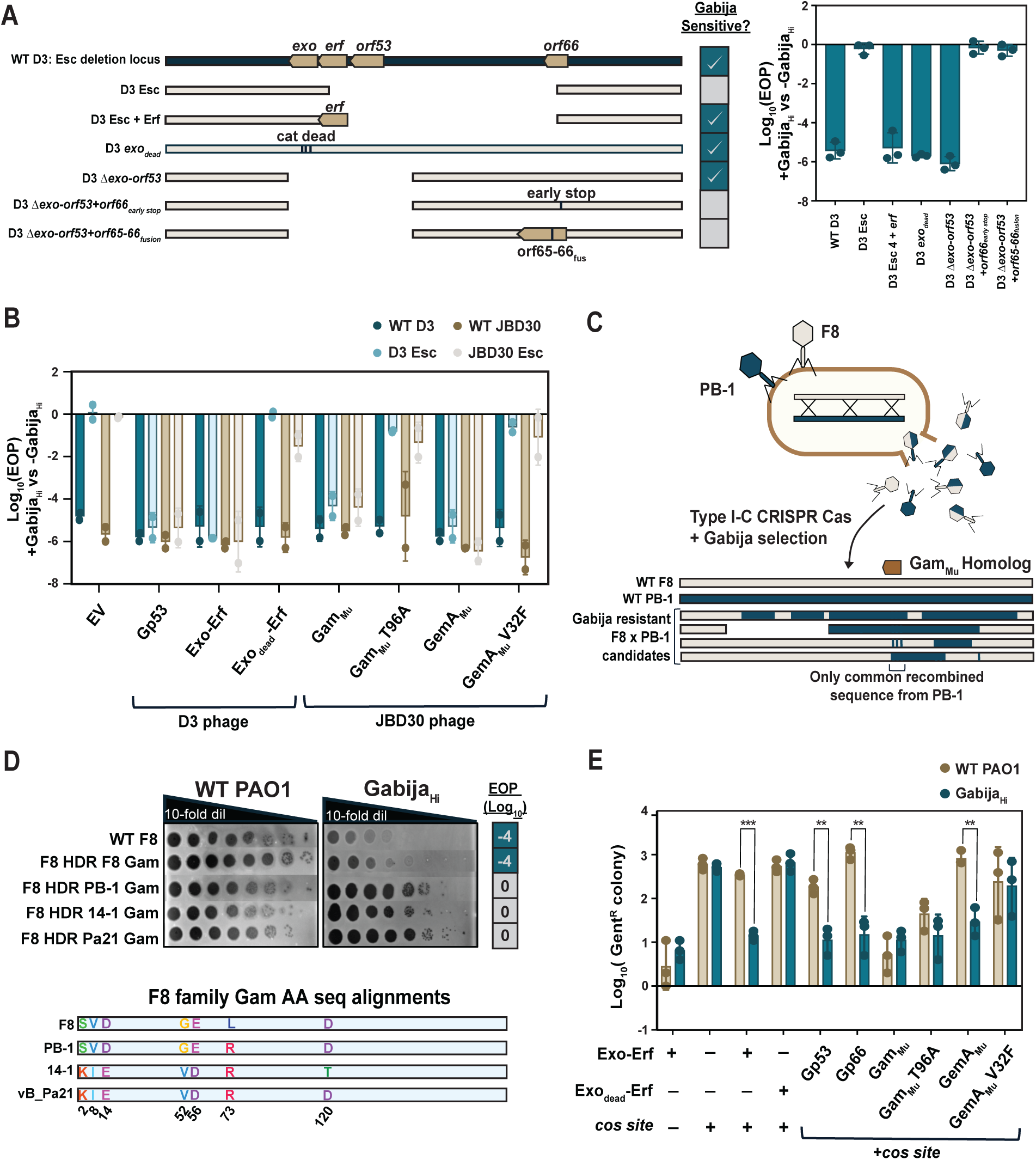
Gabija is licensed by a diverse list of phage genes and systems A. Schematic of the escape phage deletion locus of WT D3 and recombinant phages and their sensitivity to Gabija. EOP of the D3 recombinant phages is shown on the right (n=3). Data are mean ± SD. B. EOP of D3 (blues) and JBD30 (browns) phages in the presence of candidate Gabija licensing phage proteins (n=3). Data are mean ± SD. C. Schematic of F8 (tan) - PB-1 (blue) crossing experiment and example hybrid genomes. Hybrid phages were selected on a strain encoding Gabija and Type I-C CRISPR Cas targeting either *orf12* or *orf51*. A total of 6 unique hybrids were isolated and sequenced. D. Plaque assays of WT F8 and F8 recombinant Gam mutants on WT PAO1 and Gabija_Hi_. Below are amino acid sequence alignments of the Gam protein from each tested phage in the F8 family. HDR of 14-1 *gam* (Gabija sensitive) into F8 resulted in the acquisition of all diverged residues except for D120T, as recombinant phages were selected with Gabija. E. D3 transduction efficiency of *cos* plasmids encoding Gabija licensing phage proteins (log-scale of Gent^R^ colony counts) with (blue) or without (tan) Gabija (n=3). Data are mean ± SD.

To test if these three phage systems are the only Gabija licensing factors in D3, we engineered *orf51-53* deletion mutants. D3Δ*orf51-53* phages do not become resistant to Gabija, but exposure to Gabija did result in the generation of new spontaneous escape phages. Unlike WT D3 phages that must acquire large deletions to escape Gabija, these phages derived from D3Δ*orf51-53* acquired one of two unique escape mutations in *orf66* that encode either an early stop codon or a frameshift mutation that fuses orf66 to orf65. This confirms that D3 encodes three redundant Gabija licensing factors - Exo-Erf, gp53, and gp66 - and mutations or deletions in all three are necessary to achieve full Gabija escape.

While *Caudoviricetes* phages (i.e. tailed phages) inject linear DNA into the cell, not all encode a recombination module like Exo-Erf or homologs of gp53 and gp66. We therefore next wondered how many distinct phages are restricted by the same Gabija system. We first looked to the unrelated phages JBD30 and F8. From JBD30, we were able to isolate a single rare escape mutant that had three single nucleotide polymorphisms: gp13 T96A, gp20 V32F, and gp47 G215R. Complementation with either WT gp13 or gp20 restored Gabija’s ability to target the JBD30 escape phage, but WT gp47 did not, suggesting that gp13 and gp20 independently sensitize the phage to Gabija (Figure 4B). Complementation was lost when gp13 or gp20 were given their respective escape mutations.

JBD30 gp13 is homologous to Mu phage’s Gam_Mu_ (61% sequence identity), not to be confused with the DNA mimic inhibitor of RecBCD, λ Gam_λ_. Gam_Mu_ dimerizes to bind at DNA ends and blocks loading of RecBCD, a widespread prokaryotic DNA repair and recombination complex^31–33^. Gam_Mu_ is also homologous to the end-binding eukaryotic NHEJ protein Ku^31^. gp20 is homologous to Mu phage’s GemA, which is a part of the GemA-GemB *gem* operon that has not been characterized at the molecular level^34–36^.

The F8 phage family (*Pbunavirus*) had varied resistance to Gabija restriction (Figure 1B); these phages share ∼89-95% nucleotide identity, suggesting that resistance requires minimal genetic changes or phage-encoded inhibitors. Phages F8 and 14-1 are sensitive to Gabija and have high genome-wide sequence identity with either PB-1 or vB_Pa21, respectively, both of which are Gabija-resistant^37,38^. To identify the genetic determinant of Gabija targeting for these phages, we created phage recombinants by crossing F8 (Gabija-sensitive) with PB-1 (Gabija resistant) during co-infection in PAO1 followed with selection for phages resistant to Gabija and Type I-C CRISPR-Cas targeting PB-1 at one of two distinct arbitrarily chosen loci (*orf12* or *orf51*) (Figure 4C). Whole genome sequencing of hybrids that grew on both Gabija and CRISPR-Cas revealed only one PB-1 sequence that was consistently found across all hybrid phages, which spanned *orf54*. Homology searches revealed that *orf54* encodes another Gam_Mu_-like homolog. F8 Gam shares only 14.3% sequence identity with JBD30 Gam, suggesting that F8 Gam is a diverged DNA end-binding protein.

One amino acid difference was identified when comparing Gam_F8_ to Gam_PB-1_, and Gam_14-1_ to Gam_vB_Pa21_, but 5-6 substitutions exist between the two groups (Figure 4D). The Gabija resistant phages encode Gam with Arg at position 73, while the Gabija sensitive phages encode Leu at this position. Homology directed repair (HDR) - mediated insertion of the L73R substitution was sufficient to enable escape. In fact, insertion of any of the Gam homologs from a Gabija resistant phage into F8 phage led to full resistance against Gabija. These data suggest that Gam_F8_, likely through its ability to bind to the phage DNA ends, licenses Gabija targeting of phage F8. Thus, the L73R substitution, identified as a naturally circulating allele in this population, is sufficient for Gabija escape in this phage family. Interestingly, this L73 residue falls within Gam_Mu_’s leucine zipper, which suggests that the Arg substitution alters Gam_Mu_’s DNA binding affinity.

We next asked if the mechanism of Gabija activation needs to be phage specific. Interestingly, expression of D3 *exo-erf* or *orf53* led to cross-complementation of Gabija targeting against the JBD30 escape phage. This was also true for JBD30 Gam_Mu_ or JBD30 GemA_Mu_ with D3 escape phage (Figure 4B). These data suggest that end-binding proteins can license Gabija sensitivity in a non-phage-specific manner. To determine if these phage factors that license Gabija during phage infection are indeed sufficient for Gabija restriction of an injected linear dsDNA substrate, we returned to the D3 transduction assay (Figure 4E). Again, we observed a strong 1-2 log decrease in transduction of plasmids containing either D3’s *orf53* or *orf66* or JBD30’s *gemA* in Gabija_Hi_ compared to WT PAO1, suggesting that expression of any of these proteins leads to Gabija activity towards that DNA substrate. JBD30 *gam* expression, however, antagonizes transduction in a Gabija-independent manner. Together, we show that three distinct phages have 6 unique phage systems (Exo-Erf_D3_) or genes (orf53_D3_, orf66_D3_, Gam_JBD30_, GemA_JBD30_, Gam_F8_) that each independently license Gabija targeting, three of which are DNA-end binding systems.

### RecBCD absence on DNA sensitizes it to Gabija

With multiple unrelated phage genes identified as necessary and sufficient for licensing phage restriction by Gabija, we next investigated the mechanism of activation. We considered that each of these proteins directly binding and activating GajA-GajB is unlikely, particularly the proteins with DNA-binding activity. Rather, we hypothesized that Gabija activity may be a consequence of phage DNA binding by these “Gabija licensing” proteins. To test this hypothesis, we queried the involvement of a host complex that is well known to be associated with linear DNA, RecBCD. This was motivated by two reasons. First, our implication of Gam_Mu_ homologs in sensitizing phage to Gabija suggests that the local antagonism of RecBCD loading onto the phage genome may, in fact, license Gabija activity. Second, the related GajA homolog, OLD nuclease, induces cell death in a *recB-* or *recC*-strain, suggesting the potential for OLD/GajA family enzymes to interact with unrepaired exposed ends^11,39^. RecBCD is one of the most complex known multi-unit enzymes, having nine identified enzymatic activities performed across three subunits of its 330 kDa complex^40–41^. Canonically, RecBCD is a host repair system that loads onto blunt or near-blunt ends (but not resected ends) and resects by unwinding and cleaving dsDNA into short oligonucleotides. This continues until a Chi (crossover hotspot instigator) sequence (5’-GCTGGCGC-3’ for *Pseudomonas*, Chi_Pa_) is encountered, upon which the Chi site is nicked forming a 3’-OH, RecA is recruited, and recombination occurs with homologous DNA^40–42^. The Chi sequence was originally regarded to be a host-specific sequence that allowed RecBCD to act as an antiphage DNA-targeting system against invading phages. However, recent findings have shown that Chi is common across many phages^43,44^. *P. aeruginosa* RecBCD shares ∼40% amino acid identity with *E. coli* strain K12’s RecBCD.

To determine if host RecBCD participates in Gabija activity, we first generated a *recB* mutant PAO1 strain by introducing an early stop codon in the gene. In the absence of Gabija, all phages replicated on the *recB^-^* strain like they did on WT PAO1. Interestingly, a Gabija_Hi_ *recB*^-^ strain was not viable, suggesting that highly expressed Gabija, like OLD, can self-target the host genome. However, the less active Gabija_Lo_ construct (Figure 5A) did not induce self-targeting in the *recB^-^* strain and still efficiently targeted phages D3 and JBD30. Therefore, RecBCD is not required for Gabija function but is required to protect self. Surprisingly however, we observed that the D3 and JBD30 Gabija escape phages lost the ability to escape Gabija in the *recB*^-^ background and were restricted to the same level as WT phages (Figure 5B). Escape phages, therefore, resist Gabija in a RecBCD-dependent manner, suggesting that RecBCD activity protects them from Gabija. This explains why phage expression of a local RecBCD antagonist like Mu_Gam_ sensitized phage to Gabija. Exo-Erf also likely resects the near-blunt ends that RecBCD requires to load, which sensitizes WT phage to Gabija (Figure 5C).

**Figure 5.**
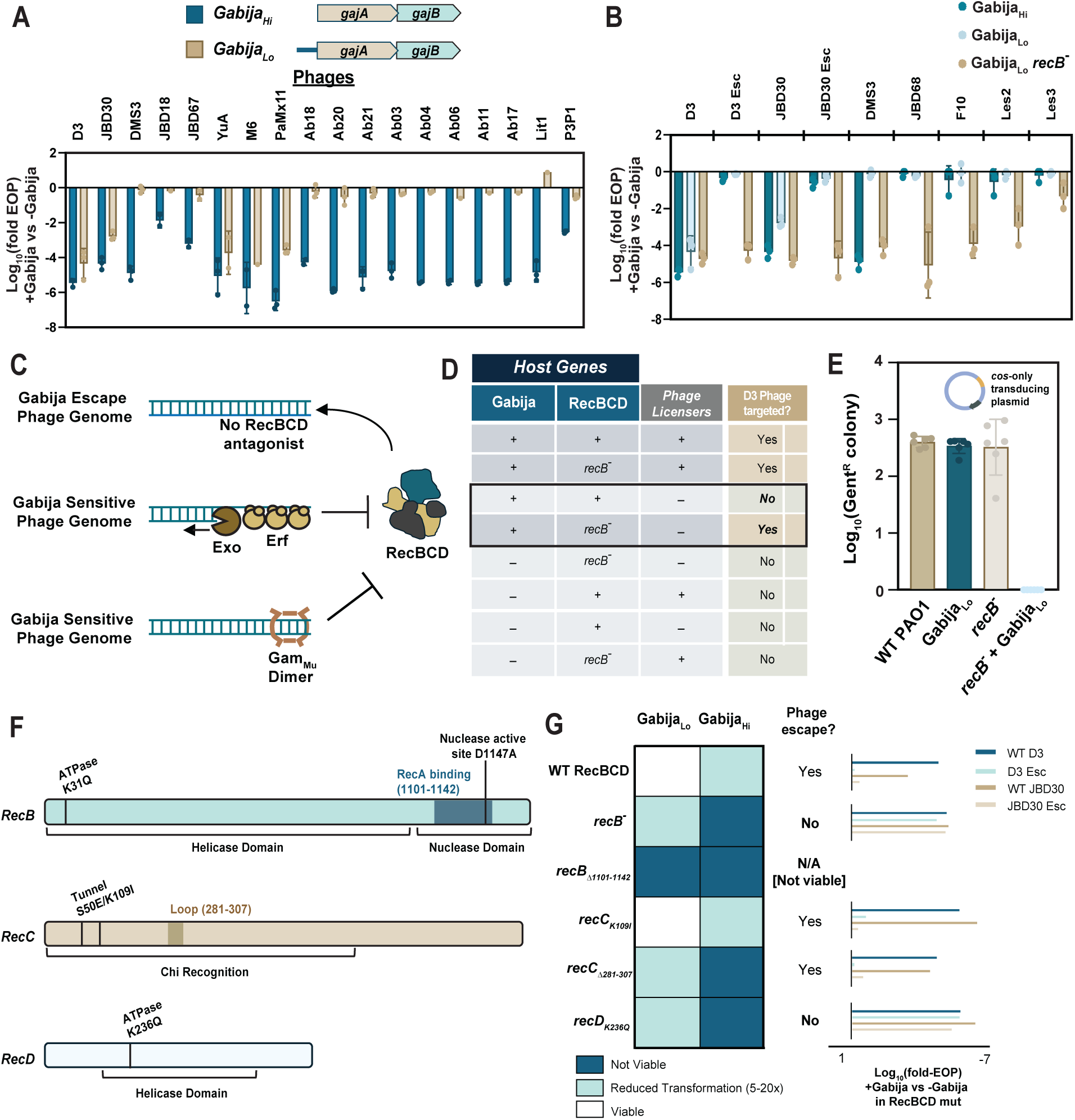
Gabija does not target DNA in the presence of RecBCD activity A. Inclusion of the native 250 bp upstream sequence (blue bar in Gabija_Lo_) of Gabija suppresses its activity of a large number of phages relative to GajA-GajB alone. Schematic of Gabija_Hi_ and Gabija_Lo_ is shown on the left. EOP of phages is shown on the right (n=3). Data are mean ± SD. B. EOP of phages on Gabija_Hi_, Gabija_Lo_, and Gabija_Lo_ in a recB^-^ strain (n=3). Data are mean ± SD. C. Schematic hypothesizing mechanism of inhibition of RecBCD loading on DNA ends. D. Table of phage sensitivity +/- Gabija, RecBCD, or Gabija licensing phage genes. The key defining genetic conditions suggesting RecBCD’s role in protecting DNA from Gabija activity are outlined in black. E. Transduction efficiency (log-scale of Gent^R^ colony counts) of the *cos*-only plasmid into strains +/- Gabija_Lo_ or RecB (n=3). Data are mean ± SD. F. Diagram of *recB, recC,* and *recD* with functional domains and tested mutations. G. Assaying loss of DNA protection upon introduction of RecBCD mutations. Heatmap indicates viability of Gabija-RecBCD mutant combinations in the absence of phage using a readout of surviving colonies when Gabija is introduced into the Tn7 site (n=2). EOP of WT and escape phages are shown on the left (n=3).

If RecBCD is indeed protecting phage DNA from Gabija, one might expect that phages that naturally do not antagonize RecBCD should resist Gabija in a RecBCD-dependent manner. Consistent with this prediction, we saw that certain phages that are normally fully or semi-resistant to Gabija such as JBD68 and DMS3, respectively, became more restricted by Gabija in a *recB*^-^ background (Figure 5B). This suggests that these phages do not exclude RecBCD efficiently and are, thus, normally protected from Gabija. Deletion of *recA,* on the other hand, had no impact on Gabija escape (Figure S4), suggesting that Gabija’s targeting mechanism is not generalized to repair pathways, but rather, specific to RecBCD loading, translocation, or nicking activities. A summary of the host and phage genetic requirements for phage targeting is in Figure 5D.

Why aren’t phages simply restricted by RecBCD? Certain lambdoid phages, such as JBD68 and F10, are rich in Chi_Pa_ sites (18 and 20, respectively). These phages may utilize RecBCD for their own purposes (i.e. some phages recruit RecBCD for circularization/packaging^44,45^), suggesting that phage genomes that adopt RecBCD for their own uses would not be targeted by Gabija. Other lambdoid phages may have fewer to no Chi sites but rely on RecBCD occlusion and expression of their own recombination machinery. D3 has 7 Chi_Pa_ sites distributed between *orf5*-*orf27*, which may explain why the escape phages are able to replicate in the presence of RecBCD despite losing *exo*-*erf*. Together, when RecBCD is non-functional, both host genomes and phages become sensitive to Gabija. This generates a prediction that any DNA end may be a target of Gabija in the recB^-^ cells. Indeed, Gabija restricted the transducing *cos*-only plasmid only in a *recB*^-^ strain, suggesting that the absence of RecBCD is sufficient to license targeting of a linear DNA substrate independently of phage licensing proteins (Figure 5E).

Lastly, we sought to understand, at a biochemical level, what mechanistic aspect of RecBCD protects escape phage and host DNA from Gabija. We generated *recB*, *recC*, or *recD* mutants in PAO1 (Figure 5F and 5G) and assayed for two indicators of alteration to Gabija activity: the loss of protection against self-targeting by measuring the relative number of surviving colonies with inserted Gabija_Hi_ or Gabija_Lo_ systems and resensitization of escape phages to Gabija_Lo_ in viable cells (Figure 5G). Strikingly, a *recD*_K236Q_ ATPase mutant is self-targeted by Gabija_Hi_ and restores Gabija_Lo_ targeting of escape phages. This suggests that RecD’s helicase activities are critical for protection of both host and escape phage DNA against Gabija. Interestingly, a *recB*_Δ1101-1142_ mutant missing the C-terminal putative RecA-recruiting/nuclease domain is viable alone, but unable to coexist with either Gabija_Hit_ or Gabija_Lo_. Due to the high Gabija-dependent toxicity, we were unable to assay for escape phage resensitization. Nonetheless, this reveals that RecB nuclease activity is also important for host DNA protection. A *recC*_K109I_ mutation in the tunnel region, which has previously been shown to reduce RecBCD’s Chi activity and high frequency recombination (Hfr)^40,46,47^, phenocopies WT PAO1, suggesting that these functions may not be critical for protection from Gabija.

This is, perhaps, consistent with the observation that a *recA* null mutation, which presumably eliminates repair/recombination, does not resensitize escape phages to Gabija. However, a *recC*_Δ281-307_ deletion mutant in the putative RecC loop region, which is required for full Chi activity in *E. coli*, partially phenocopies our original *recB*^-^’s toxicity profile, with reduced insertion rates of Gabija_Lo_ and complete non-viability with Gabija_Hi_ insertions. Interestingly, escape phages do not become targeted in this mutant, suggesting that this region alone is not solely responsible for escape phage protection. Together, we show that RecBCD functions, particularly RecB and RecD translocation and/or nuclease activities, are crucial for protecting self and escape phage DNA against Gabija’s DNA-targeting activity and, thus, is responsible for Gabija’s self vs non-self substrate determination. The translocation and nicking of DNA by RecBCD thus likely modifies the DNA substrate to render it Gabija resistant, explaining why phages that block RecBCD loading have Gabija-sensitive ends.

## Discussion

In this study, we report the phage and host determinants required for Gabija activation *in vivo*, highlighting contradicting evidence against the currently proposed abortive nucleotide depletion activation mechanism for Gabija^16,17^. Namely, we discovered that Gabija_PA14_ restricts D3 phage genome circularization, without any observed cellular growth arrest, and limits transduction of a phage-injected DNA substrate. Moreover, DNA translocated, unwound, and/or nicked by RecBCD are Gabija resistant, but exclusion or ablation of RecBCD activity leads to Gabija activity that prevents end-joining (Figure 5). Together, these data implicate a battle for linear DNA between phage proteins, Gabija, and a conserved host repair complex. Accordingly, Gabija sensitivity can be bypassed by inducing prophages, which proceed directly to the circular form without an exposed linear intermediate. This result additionally supports that phage DNA replication per se is not detected nor interfered with by Gabija.

*Caudoviricetes* genomes are injected in linear form and use myriad mechanisms to proceed to circularization^48–54^. DNA ends are therefore an apposite hotspot for host-pathogen molecular interactions and, more specifically, an opportunistic battleground for DNA-targeting systems to monitor and act on. For example, the DNA end-sensing Shedu complex directly binds to and nicks linear dsDNA^55,56^. Furthermore, many bacteria encode RecBCD and similar systems that load onto DNA ends to enact processes towards repair, which may interfere with various phage linear DNA intermediates. Therefore, some phages will exclude or inhibit RecBCD. Indeed, it has been suggested that Retron Ec48 and OLD respond to RecBCD antagonism with lambda phages escaping OLD by deleting their entire red recombination system (Figure 6B)^7,10,13,57^. However, RecBCD is not necessarily anti-phage^43^, as other Chi-abundant phages rely on RecBCD binding and nicking for assistance with circularization, replication mechanisms (i.e. shifting phages from theta to rolling circle replication), or even packaging^26,40,43,44^. We further showed that RecBCD nicking and translocation activities alter linear dsDNA to an undesirable substrate for Gabija, perhaps by complex translocation, DNA oligo generation, or ssDNA loop formation^40^. Interestingly, across sequenced prokaryotic genomes, Gabija is not strictly correlated with RecBCD, however, analogous repair systems including AddAB and AdnAB are widespread, and thus, we reason that phages are likely commonly equipped with antagonists towards these systems^58,59^. This may explain why many distinct phage families are targeted by Gabija.

**Figure 6.**
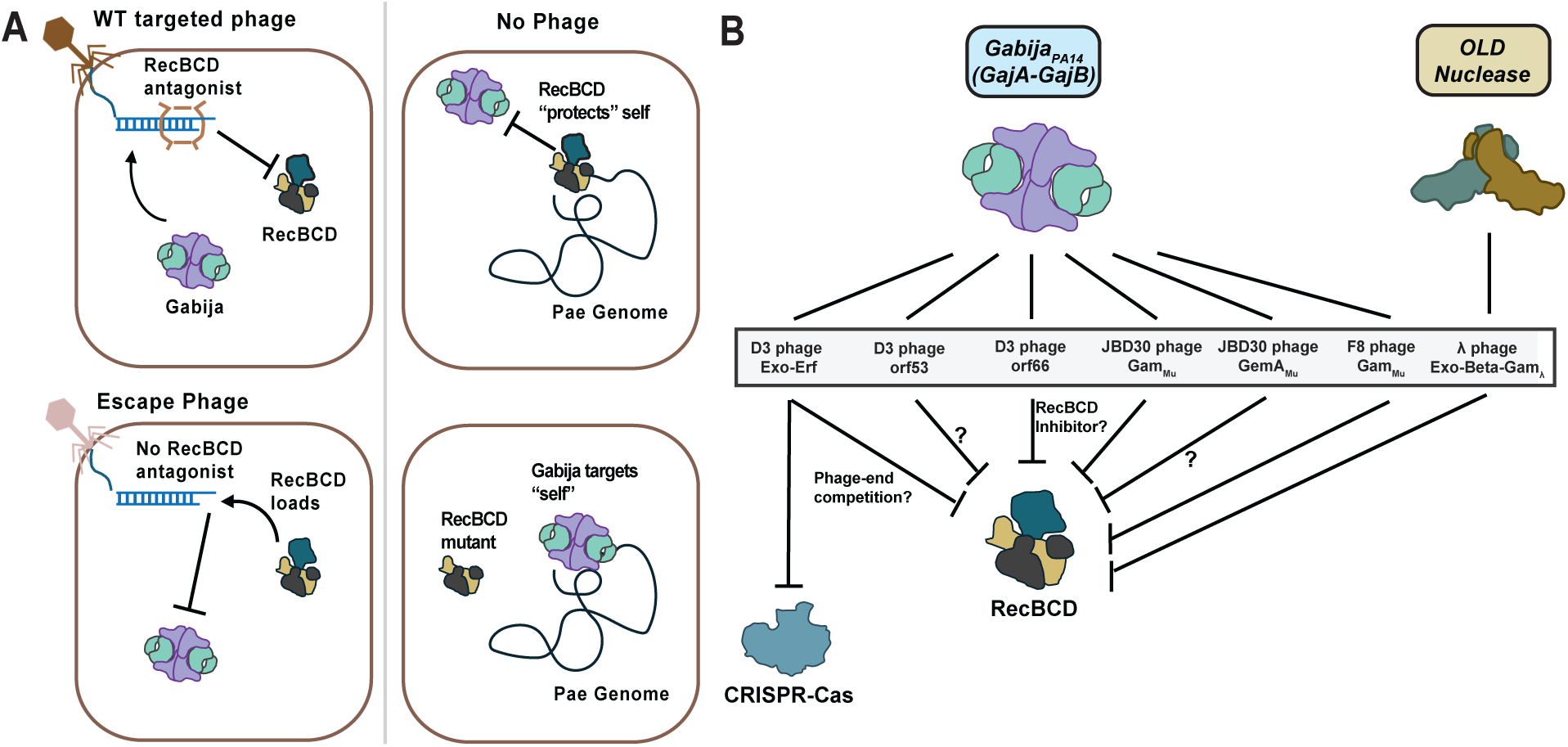
Graphical summary of the Gabija mechanism A. Gabija targets DNA that is not acted upon by RecBCD B. A summary of the phage proteins that license Gabija targeting, highlighting parallels found in GajA homolog OLD and interplay with CRISPR-Cas systems.

The D3 escape phages generated remarkably massive - up to 16.9 kb - spontaneous deletion mutations in seemingly nonessential regions. Such a concept of readily removable genomic regions would be an interesting mechanism for phages and other pathogenic MGEs to exchange and harness defense evasion genes. Akin to appendage autotomy, such regions can be acquired and discarded upon encountering defense systems that might target or detect these regions. This suggests that escape from systems that detect multiple genes of a phage may not always be difficult but can induce a trade-off as observed here with CRISPR-Cas targeting. This observation presents a unique example of how phages respond to multiple layers of host defense, particularly against two DNA-targeting systems that comprise a considerable portion of the known bacterial antiphage arsenal.

Together, we propose that Gabija interferes with linear DNA that is devoid of RecBCD interaction as a proxy indicator of non-self substrates (Figure 6A). Phages that are equipped with RecBCD antagonists prevent RecBCD loading on the phage gDNA by blocking it directly (Gam Mu) or resecting the ends (Exo-Erf). Perhaps, phage antagonization of RecBCD normally prevents this host complex from interfering with replication or repair intermediates, explaining why MGEs encode mechanisms to not only block RecBCD, but execute repair and recombination independently. RecBCD antagonism is then an appropriate invasion alarm that Gabija recognizes as local absence of RecBCD function on phage gDNA. Conversely, phages that allow RecBCD loading and translocation become “protected” from Gabija targeting. In a WT host genome, RecBCD also loads onto and repairs linearized host DNA. This normally protects host gDNA from Gabija self-targeting when there is DNA damage. However, if RecBCD is mutated or directly inhibited, global RecBCD function is abrogated, allowing Gabija to prevent circularization of on “unprotected” DNA regardless of whether it is self- or non-self-DNA. Importantly, despite this potential to target self upon prolonged RecBCD inactivation, all available data demonstrate that self-DNA is not the target during *bona fide* phage immunity.

## Methods

### General Conditions

All experiments assaying Gabija and RecBCD activity were performed at 30 °C unless otherwise specified. All plasmids were sequenced by primers flanking both ends of the multiple cloning site (Quintara) or whole plasmid sequenced (Plasmidsaurus). All DNA concentrations were measured using the dsDNA High Sensitivity Qubit kit (Thermo Fisher Scientific).

### Bacterial strains, phages, and plasmids

The key resources table includes all bacterial and phage strains used in this study. Bacterial strains were cultured in lysogeny broth (LB) at 37 °C with aeration at 175 rpm. Assays involving Gabija antiphage activity were carried out at 30 °C. *P. aeruginosa* plasmids were maintained by 50 ug ml^-1^ Gentamycin (15 ug ml^-1^ for *E. coli*). Bacteria were plated on LB agar, with 10 mM MgSO_4_ included for phage assays. When applicable, attTn7-inserted gene expression was induced with 1 mM IPTG while pHERD30T gene expression was induced with 0.1% L-arabinose. Cloning plasmids were transformed into XL1-Blue or DH5ɑ competent cells and selected on [15 mg/mL] gentamicin.

Phage lysates were made by adding 10 ul of high titer (10^7–9^ pfu) phages to 150 ul of overnight PAO1 cultures (or PAO1-Gabija^PA14^ cultures for escaper phages) and allowing adsorption for up to 20 minutes. The infected cultures were then plated in 0.7% top agar with 10 mM MgSO_4_ and allowed to grow overnight at 30 °C. SM buffer was then added to plates and incubated at RT for at least 10 minutes, and then transferred into microfuge tubes with 2% chloroform. Lysate mixtures were pelleted, and supernatants were transferred to fresh tubes with 2% chloroform then stored at 4 °C. The key resources table also includes detailed plasmid and oligo information.

### Gabija and RecBCD strains

The GajA-GajB system was amplified from *P. aeruginosa* strain PA14 and cloned into pHERD30T for episomal expression or pUC18-mini-Tn7T-LAC for chromosomal integration (see below). Two constructs for GajA-GajB in mini-Tn7 were generated, one with just GajA-GajB and one including 250 bp native sequence flanking either side of Gabija. mini-Tn7 insertions were generated as previously described and gent^R^ were removed^60,61^. GajA and GajB mutants were generated by PCR amplification of native sequences using primers with respective SNPs. VD045 (AHET01000033.1) GajA-GajB sequences were codon optimized for *P. aeruginosa* and ordered from TWIST Biosciences as two gBlocks with matching internal overhangs and external overhangs with pHERD30T multiple cloning site and inserted into pHERD30T with Hi-Fi DNA Gibson assembly (NEB). The *recB^-^* mutant was generated by Balint Csorgo using ssDNA-mediated recombineering via pORTMAGE-Pa1 plasmid^62^. The RecBCD point and deletion mutants were generated by two-step allelic exchange as previously described (see below)^63^.

### Lysogen formation

Lysogens were isolated by first streaking a high titer clearing of the respective phage in PAO1 on LB agar. Single colonies were patch plated and tested for superinfection exclusion via O-antigen serotype conversion by spot assays^64,65^. Candidates that lacked plaques were assayed for spontaneous prophage induction by spot titration assay with overnight supernatant.

### Chromosomal integration of Gabija constructs

PA14 GajA-GajB constructs (+/- 250 bp upstream native sequence) were amplified from *P. aeruginosa* strain PA14 WT and inserted into Sac1-Pst1 linearized pUC18-mini-Tn7T-LAC vector by Hi-Fi DNA Gibson assembly (NEB). Plasmids were transformed into NEB DH5⍺ competent cells. For chromosomal insertion, miniTn7 constructs (including empty vector controls) were transformed along with the transposase helper vector pTNS3 into PAO1 and selected on gentamicin LB agar plates. Candidate integrants were screened by colony PCR using primers PTn7R and PglmS-down coupled with a second primer set that paired PglmS-down with an integrant gene-specific primer. Integrants were confirmed by sequencing with primers flanking the Tn7 integration site. FLP-recombinase-mediated gentamicin excision was performed as previously described with pFLP2 vector^61^.

### Plasmid construction and transformations

For complementation and expression assays, *P. aeruginosa* and phage gene constructs were either synthesized via PCR or ordered as gBlocks from TWIST Biosciences. *E. coli* constructs were codon optimized for *P. aeruginosa* and ordered as gBlocks from TWIST Biosciences. Gabija anti-defense gene sequences were provided by Rotem Sorek and Phillip Kranzusch.

### Plaque assays

10-fold serial dilutions were made with phage lysate stocks in SM buffer. These were then spotted in 2 uL aliquots onto bacterial lawns consisting of 150 ul of overnight culture in 3 mL of 0.7% top agar with 10 mM MgSO_4_ poured onto LB agar plates in 1.5 cm circular plates (300 ul of overnight culture in 6 mL top agar for larger rectangular plates) and allowed to dry next to an open flame for ∼5 min, then incubated overnight at 30 °C.

### Phage and bacterial genomic DNA isolation and purification

An equal volume of lysis buffer (final concentration 10 mM Tris pH 7.5, 1 mM EDTA, 100 µg ml^−1^ Proteinase K, 100 µg ml^−1^ RNaseA and 0.5% SDS) was added to 300 ul of bacterial overnight culture or 400 ul of high titer phage sample. Samples were incubated in Eppendorf Thermomixer Cs at 37 °C for 30 minutes then 50 °C for 1 hour shaking at 500 rpm. Genomic DNA was then isolated with Genomic DNA Clean and Concentrator kits (Zymo Research).

### Whole genome sequencing and analysis

For both phage and bacterial genome sequencing, 20-100 ng of gDNA were tagmented with Illumina’s DNA Prep kit and amplified with Phusion polymerase using Illumina Nextera compatible primers designed by Sukrit Silas. Libraries were then pooled to ensure that coverage of bacterial genomes retained at least 100 reads per 300-400 bp. Demultiplexing and alignments were then generated with scripts generated in the Bondy-Denomy lab. Vcfs of single nucleotide polymorphisms and indels were generated by breseq (Barrick Lab). *de novo* assembly of genomes was generated with Spades (Bankevich et al).

Genomic DNA (50–100 ng) was used to prepare whole-genome-sequencing libraries using the Illumina DNA Prep Kit (previously, Nextera Flex Library Prep Kit). A modified protocol was used, with fivefold reduced quantities of tagmentation reagents per preparation, except for the bead-washing step, in which the recommended 100 µl of tagment wash buffer was used. On-bead PCR indexing amplification was performed using custom-ordered indexing primers (IDT) matching the Illumina Nextera Index Kit sequences and 2× Phusion Master Mix (NEB). PCR reactions were amplified for 9–11 cycles, and subsequently resolved by agarose gel electrophoresis. DNA products were excised around the 400-base pair size range and purified using the Zymo Gel DNA Recovery kit (Zymo Research). Libraries were quantified by Qubit. Libraries were pooled in equimolar ratios and sequenced on Illumina MiSeq using 150 cycle v.3 reagents (single end: read 1, 150 cycles; index 1, eight cycles; index 2, eight cycles. Paired end: read 1, 75 cycles; index 1, eight cycles; index 2, eight cycles; read 2, 75 cycles). Data were demultiplexed on instrument and trimmed using cutadapt (v.1.15) to remove Nextera adaptors. Trimmed reads were mapped using Bowtie 2.0 (ref. 24) (– very-sensitive-local alignments) and alignments visualized using IGV (v.2.11.0). Mutations were called if present in over 90% of sequencing reads at loci with at least 20× coverage.

### Phage Transduction Experiments

The D3 *cos* sequence and flanking upstream and downstream 250 bp sequences were cloned into the non-coding region of pHERD30T downstream of the aacC1 gene. Flanking regions included putative terminase binding sites. Subsequently, WT or mutant phage genes were inserted into the pHERD30T multiple cloning site via Gibson assembly. Full plate lysates of WT D3 or escape D3 phages grown in PAO1 harboring each of these plasmids were isolated, then treated with DNase I (NEB). Pa strains were infected with each lysate at an MOI of 0.1-2, allowed to adsorb for ∼15-20 minutes, plated on Gent50 0.1% arabinose 10mM MgSO_4_ LB agar, and incubated overnight at 30 °C. Colonies were manually counted the following day.

### Phage recombineering

Recombinant vectors were generated to have desired mutations flanked by 500 upstream and downstream homology arms. Phage lysates were generated on PAO1 with recombinant vectors. Recombinant phages were then selected for by Type I-C CRIPSR-Cas targeting with guides against the WT sequence at mutation sites or Gabija. When generating the add-back recombinant D3 escaper phages, AcrIF2/IC2 was included within the homology arm cassette. Recombinant phages were isolated by a two-step CRISPR-Cas3 selection first with a guide against the insertion site followed by using a guide against an essential D3 early gene for robust selection.

### Two-step allelic exchange

TWIST Bioscience gBlocks of a desired mutation flanked on either side by 500 bp homology arms of the landing site were cloned via Gibson into the multiple cloning site of allelic exchange plasmid pMQ30. These plasmids were then electroporated into SM10 *E. coli* cells and selected for on gentamicin followed by conjugation with PAO1 or PA14 by cross streaking and allowed to grow at 37 °C for 24+ hours depending on growth rate. Recombinant Pa strains were then selected for on Vogel-Bonner minimal medium (VBMM) agar with 50 mg/mL gentamicin. Gentamicin resistance was removed by selection on no-salt LB 15% sucrose LB agar^63^. Mutations were screened by PCR followed by next generation sequencing (Illumina Nextera kits).

### Single step infections

Phage infection of log phase (1:100 dilution of overnight cultures grown to ∼0.5 OD and concentrated 2X) cultures were carried out at an MOI of 1 in 10 mM MgSO_4_ LB media. An adsorption period of 5 or 20 minutes was followed by a wash step. Infections were then diluted 2X and incubated in a 30 °C shaker bath. To measure timing of phage burst, 50 uL of supernatant was then added to 450 uL of SM buffer at subsequent time points. Samples were titered by plaque assays. To extract gDNA, 2 mL aliquots of infected cultures were flash frozen with liquid nitrogen. Samples were then thawed on ice, pelleted and resuspended in 300 ul of SM Buffer. gDNA was then extracted as described above.

### Southern Blotting

1-2 ug of each gDNA sample (same ug across each set of infections) was digested with BstBI at 65 °C for 3 hours. Digested samples were run on 0.8% TAE agarose gels at 60V 300 amp for 4.25 hours in Thermofisher OWL B1 mini EasyCast^TM^ rigs. Gels were then blotted for 2 hours with 0.4N NaOH 0.6M NaCl denaturing buffer followed by a 2x saline-sodium citrate (SSC) solution overnight onto Amersham Hybond-N^+^ membranes. Membranes were cross-linked in a Stratagene UV Stratalinker then prehybridized with 100 ug ml^-1^ boiled herring sperm DNA (hsDNA) in 50% formamide, 5x Denhardt’s solution, 0.5% SDS, 6x SSC buffer for 2 hours at 42 °C. Hybridization was carried out overnight at 42 °C with P^32^-dCTP probes in the same prehybridization buffer omitting hsDNA. Whole genome probes were generated by random hexamer priming of Klenow fragment on WT D3 phage gDNA, while site-specific probes were generated by site-specific primers and gel extracted PCR products of the desired template. Membranes were then washed twice with 2x SSC 1% SDS buffer for 10 minutes at room temperature, twice for 30 minutes at 65 °C, and once for 10 minutes with 0.2x SSC 0.1% SDS buffer at room temperature before exposing on a phosphoscreen in saran wrap overnight. Images were developed on a Amersham Typhoon biomolecular imager at 50 uM resolution and analyzed in ImageQuant TL software.

### Growth assays

To analyze the growth dynamics of *P.* aeruginosa strains, overnight cultures were diluted 1:100 with 10 mM MgSO_4_ LB and respective inducers and antibiotics. For phage infection assays, a 10-fold serial dilution of phage in SM buffer was added for resulting range of MOIs from 0.0001 to 100. OD_600_ reads were collected with Agilent Biotek Gen 5 plate readers under fast orbital shaking conditions at 30 °C.

### Phylogenetic analyses

A 3x iteration PsiBLAST search in the non-redundant protein database was conducted with parameters: 70% query cover, sort by query cover, E-value of 0.005, BLOSUM62 matrix, 5000 max target sequences. Hits were then clustered via mmseqs with a minimum sequence identity of 95% and displayed as a phylogenetic tree using iTOL: Interactive Tree of Life.

### Protein structure predictions and analyses

Protein structures were predicted with AlphaFold3. Isoelectric points were predicted with Expasy’s compute pI/Mw tool. Predicted protein structure analyses were performed on ChimeraX.

## Acknowledgements

This work in the Bondy-Denomy lab was supported by the Kleberg Foundation, National Institutes of Health (NIH) (R01AI167412, J.B.D.), and National Institute of General Medical Sciences (NIGMS) (R35GM118120, G.R.S.). We thank Dr. Alan Davidson and Dr. Daan Swarts for sharing expertise on bacteriophage biology and DNA repair systems. We thank Bondy-Denomy lab members for their constructive comments.

## Author Contributions

A.H. conceived the project, designed and performed experiments, and wrote the manuscript. M.L. isolated the JBD30 escape phage and characterized Gam_Mu_ and GemA_Mu_. A.T. performed single step infections with D3 phage. A.T. assisted with the EOP phage panels. G.R. provided experimental and RecBCD advice. J.B.-D. supervised the project, designed experiments, and wrote the manuscript.

