## Supplemental Figures for "Gabija restricts phages that antagonize a conserved host DNA repair complex"

**Supplemental Data**

**Figure S1** Gabija significantly improves in phage restriction at 30ºC compared to 37ºC.

**Figure S2** Complementation of Gabija's escape phage targeting with *orf53-65*. A $\text{Gabija}_{Hi}$ strain containing *orf66* plasmid is not viable.

**Figure S3** Complementation of Gabija's escape phage targeting with Erf or Exo-Erf.

**Figure S4** Gabija does not restrict escape phages in PAO1::*recA*$^-$.

**Table S1** Predicted domains or homologs of D3 *orfs* in escaper deletions.

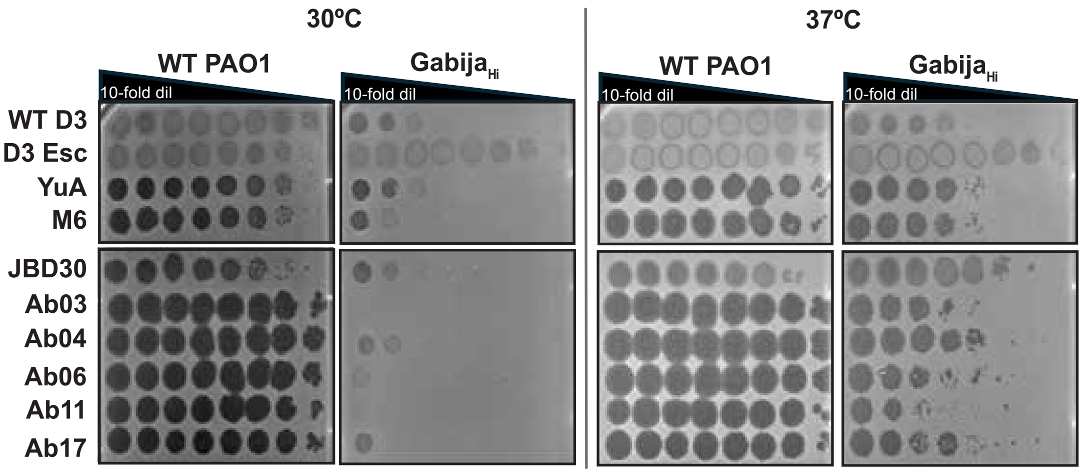

30ºC
37ºC
WT PAO1
Gabija$_{Hi}$
WT PAO1
Gabija$_{Hi}$
10-fold dil
10-fold dil
10-fold dil
10-fold dil
WT D3
D3 Esc
YuA
M6
JBD30
Ab03
Ab04
Ab06
Ab11
Ab17

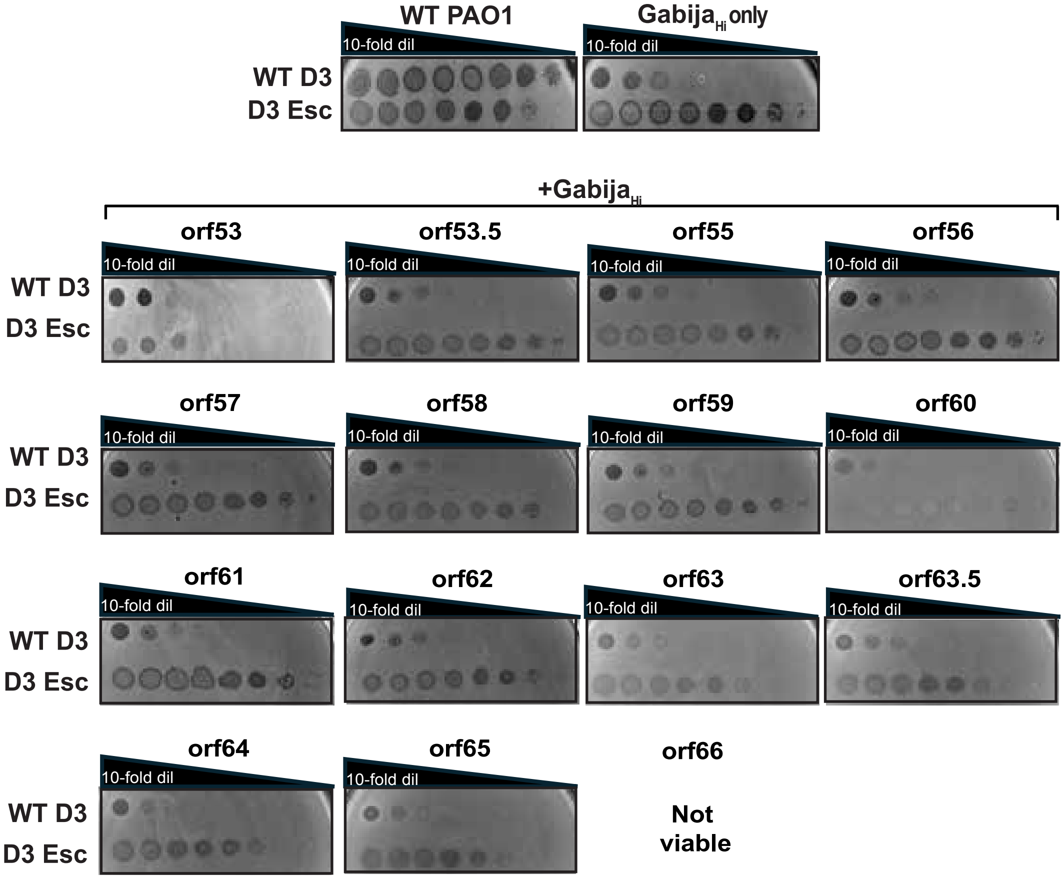

WT PAO1
Gabija$_{Hi}$ only
10-fold dil
WT D3
D3 Esc
+Gabija$_{Hi}$
orf53
orf53.5
orf55
orf56
orf57
orf58
orf59
orf60
orf61
orf62
orf63
orf63.5
orf64
orf65
orf66
Not viable

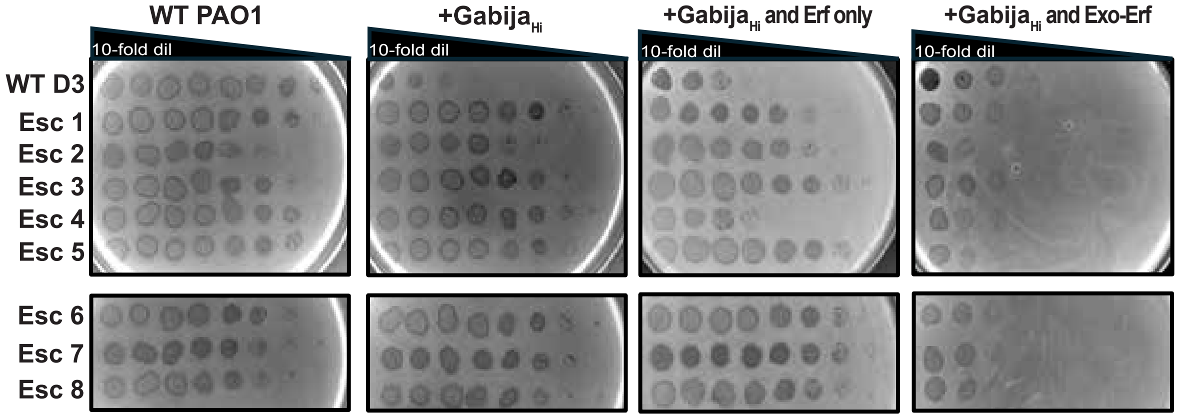
WT PAO1
+Gabija$_{Hi}$
+Gabija$_{Hi}$ and Erf only
+Gabija$_{Hi}$ and Exo-Erf
10-fold dil
10-fold dil
10-fold dil
10-fold dil
WT D3
Esc 1
Esc 2
Esc 3
Esc 4
Esc 5
Esc 6
Esc 7
Esc 8

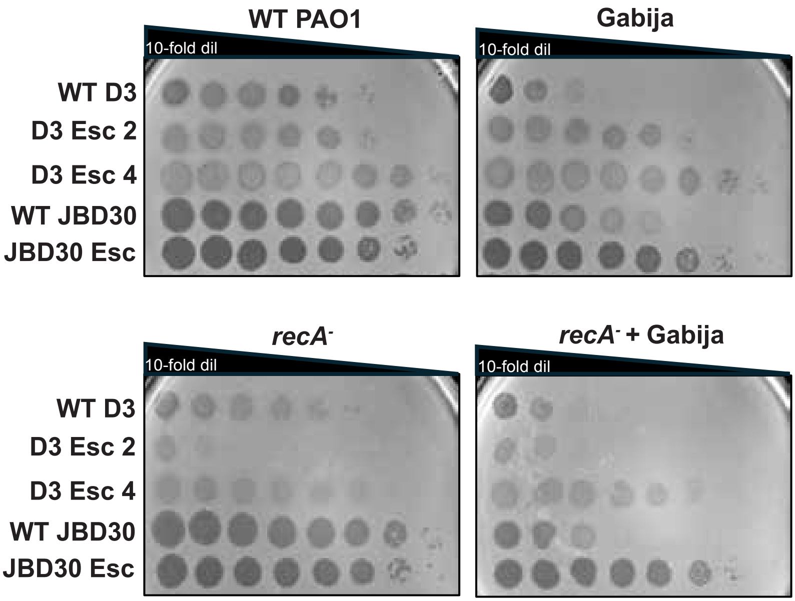
WT PAO1
Gabija
10-fold dil
WT D3
D3 Esc 2
D3 Esc 4
WT JBD30
JBD30 Esc
recA-
recA- + Gabija
